# aiMeRA: A generic modular response analysis R package and its application to estrogen and retinoic acid receptors crosstalk

**DOI:** 10.1101/2020.01.30.925800

**Authors:** Gabriel Jimenez-Dominguez, Patrice Ravel, Stéphan Jalaguier, Vincent Cavaillès, Jacques Colinge

## Abstract

Modular response analysis (MRA) is a widely used modeling technique to uncover coupling strengths in molecular networks under a steady-state condition by means of perturbation experiments. We propose an extension of this methodology to search genomic data for new associations with a network modeled by MRA and to improve the predictive accuracy of MRA models. These extensions are illustrated by exploring the cross talk between estrogen and retinoic acid receptors, two nuclear receptors implicated in several hormone-driven cancers such as breast. We also present a novel, rigorous and elegant mathematical derivation of MRA equations, which is the foundation of this work and of an R package that is freely available at https://github.com/bioinfo-ircm/aiMeRA/. This mathematical analysis should facilitate MRA understanding by newcomers.

**Author summary:** Estrogen and retinoic acid receptors play an important role in several hormone-driven cancers and share co-regulators and co-repressors that modulate their transcription factor activity. The literature shows evidence for crosstalk between these two receptors and suggests that spatial competition on the promoters could be a mechanism. We used MRA to explore the possibility that key co-repressors, i.e., NRIP1 (RIP140) and LCoR could also mediate crosstalk by exploiting new quantitative (qPCR) and RNA sequencing data. The transcription factor role of the receptors and the availability of genome-wide data enabled us to explore extensions of the MRA methodology to explore genome-wide data sets *a posteriori*, searching for genes associated with a molecular network that was sampled by perturbation experiments. Despite nearly two decades of use, we felt that MRA lacked a systematic mathematical derivation. We present here an elegant and rather simple analysis that should greatly facilitate newcomers’ understanding of MRA details. Moreover, an easy-to-use R package is released that should make MRA accessible to biology labs without mathematical expertise. Quantitative data are embedded in the R package and RNA sequencing data are available from GEO.

## Introduction

Modular response analysis (MRA) was introduced to infer the coupling between components of a biological system in a steady-state [1]. It can be applied to components at different levels of details, e.g., individual genes or subsystems such as pathways or processes. It relies on the perturbation of individual components, the so-called modules. Various developments of MRA and related methods were recently reviewed [2] but, despite its success, MRA mathematical derivation was not provided in a systematic and rigorous manner. We thus reasoned that such an analysis was needed and it would facilitate the understanding of the methodology for newcomers. It is presented as a result and is the basis of the development of an open source R library (aiMeRA) that should make MRA accessible to biology labs without mathematical expertise. We illustrate the use of the aiMeRA package by investigating the crosstalk between nuclear receptors (NRs) in a breast cancer (BC) cell line. A new extension of the method is also introduced to perform inferences at the genomic scale. The Blüthgen Lab recently released another R package to perform MRA computations [3], although with a specific focus on their particular edge-pruning and associated maximum likelihood extension of MRA [4] that is not our interest in this study.

Estrogen receptors (ERs) belong to the NR superfamily, which act as transcription factors activated upon ligand binding. The two isoforms of ERs (ERα and ERβ) are involved in the control of cell proliferation and exhibit essential functions in tissue development and homeostasis, in particular in organs related to reproduction [5]. ERα overexpression is frequently observed in breast, ovarian, endometrial, and other hormone-driven tumors. The transcriptional activity of ERs is modulated by several coregulatory complexes including coactivators and corepressors [5]. In the presence of estrogens or any agonist ligand, ERs interact preferentially with coactivators, or with a specific subclass of corepressors including nuclear receptor-interacting protein 1 (NRIP1 or RIP140) and Ligand-dependent corepressor (LCoR). NRIP1 is a corepressor of particular interest because its expression is directly induced by estrogen, i.e., NRIP1 installs a negative feedback loop to keep ER signaling under control [6]. NRIP1 abnormal expression is indeed observed in ER-driven tumors [7,8]. LCoR represses transcription of estrogen-induced gene expression [9], and NRIP1 expression was shown to be necessary for LCoR inhibitory activity in BC cells [10].

Interestingly, NRIP1 and LCoR function as corepressors for several liganded NRs. For instance, LCoR can repress vitamin D receptor (VDR), retinoic acid receptors (RARs), and RXR ligand-dependent transcription [9] in addition to ERs. Moreover, NRIP1 is a known direct target and negative regulator of RAR transcription [11].

There is experimental evidence of crosstalk between ER and RAR signaling [12]. For instance, ERα can suppress the basal expression of retinoic acid (RA)-responsive gene RARβ2, but also turns out to be necessary for its RA induction [13]. It was also found that ERα activates RARα1 expression in BC cells [14]. Other authors intersected RAR targets identified by ChIP-seq with ER binding sites to discover a significant overlap [15]. This work suggested a space competition mechanism for estrogen and RA signaling in BC. A potential cooperative interaction between RARα and ER was also shown in BC [16]. Since NRIP1 and LCoR expression can be both regulated by RAR and ER transcription, we can further hypothesize that these molecules mediate part of ER-RAR crosstalk. The induced expression of NRIP1 and LCoR by one receptor produces molecules able to repress signaling of both receptors subsequently.

We aimed at characterizing the ER-RAR-NRIP1-LCoR network, at the transcriptional level, utilizing transcript abundance measurements and MRA. Accordingly, we considered a steady-state situation in a BC (MCF7-derived) cell line that would model BC cells with or without constant estrogenic stimulation. Perturbation experiments were realized to generate quantitative PCR data unraveling interaction strengths in the network, i.e., coupling according to MRA principles.

Given the nature of ER and RAR, i.e., transcription factors, and the general ability of MRA to perform predictions [4], we introduced an extension of the method to perform genome-wide inferences exploiting mRNA sequencing (RNA-seq) data. For this purpose, we acquired whole transcriptomes under perturbed conditions identical to those used for qPCR. We first established that MRA could produce accurate results from RNA-seq data. Next, we asked whether the ER-RAR-NRIP1-LCoR network inferred by MRA could predict the mRNA abundance of estrogen-targeted genes better than a trivial model. This extension of MRA, where one or several modules do not experience perturbations, was called *unidirectional* to underline the implied absence of potential influence on the other modules.

## Results

### Mathematical derivation

MRA original paper [1] introduced the concept of modeling interdependencies (coupling) within a biological system modularly. That is, subsystems involving molecules and their relationship at a detailed level, which would not be the interest of the study, could be captured as a single module with one measurable quantity defining the overall module activity. For instance, in the case of ERα signaling, it is possible to represent the complex process of ligand-binding and transcriptional activity by a single module (Fig. 1). The activity of this module is measured by a reporter gene, which is the luciferase in MELN cells. NRIP1 and LCoR activities were determined by their respective mRNA abundances. We ask the question of the direct dependence of each module activity with respect to the other modules activity. That is, we want to compute (signed) weights to put on the directed edges of Fig. 1. The answer is searched in a steady-state through successive elementary perturbations of each module activity. Depending on the application, this framework can be applied to different molecular species and processes, e.g., protein or metabolite concentrations, protein phosphorylation levels, etc. [2].

**Fig. 1.**
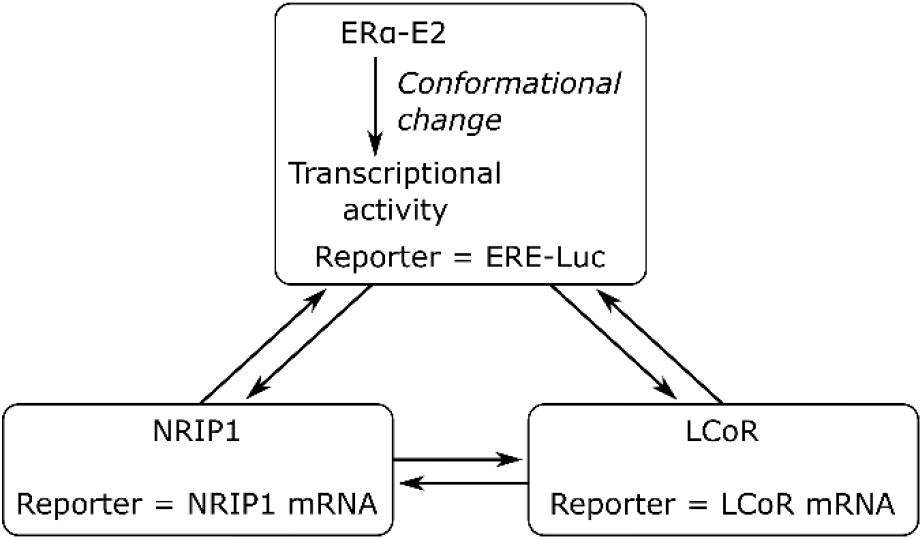
MRA general principle. Illustrated with the ERα-NRIP1-LCoR transcriptional network. Each module’s activity level is given by a measured reporter. Coupling (edge weights) is determined from perturbation experiments.

Now, in full generality, we assume that there are *n* modules whose activities are given by 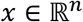. We further admit the existence of *n* intrinsic parameters, 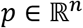, one *per* module, which are perturbed by the elementary perturbations. One can imagine mRNA abundance parameters for perturbations such as siNRIP1 or siLCoR, and numbers of available ERα-E2 bound complexes for the E2 perturbation. In other circumstances, perturbations could change affinity constants or other physical characteristics. Finally, we assume that there exist 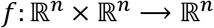 of class 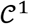 (continuously differentiable) such that

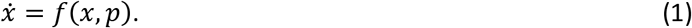

We do not need to know *f*(*x, p*) = (*f*_1_(*x, p*),…,*f_n_*(*x, p*))^*t*^ explicitly, but we need one more hypothesis that is the existence of a time *T* > 0 such that all the solutions we consider for any *p* and initial conditions of *x*, have reached a steady-state, i.e.,

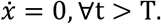

The unperturbed, basal state of the modules is denoted 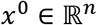 and it has corresponding parameters 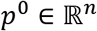. According to our hypotheses, *f*(*x*^0^,*p*^0^) = ⇔ *f_i_*(*x*^0^,*p*^0^) = 0, ∀*i* ∈ {1,…,*n*}. By the implicit function theorem, ∀*i*, there exists open neighborhoods 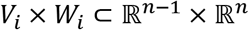 of 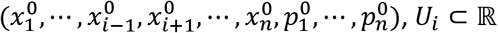 of 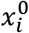, and *g_i_*:*V_i_ × W_i_* → *U_i_* (also 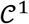) with

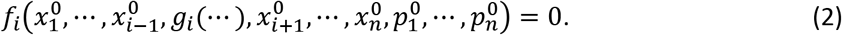

We denote *x*(*p*^0^ + Δ*p*), the steady-state corresponding to the changed parameters *p*^0^ + Δ*p*. We introduce the notation *x_j≠i_* to denote all the *x_j_* but *x_i_*. Now, if we assume that (*x_j≠i_*(*p*^0^ + Δ*p*), *p*^0^ + Δ*p*) belong to *V_i_ × W_i_* for all the perturbations considered experimentally, then by Taylor’s Formula

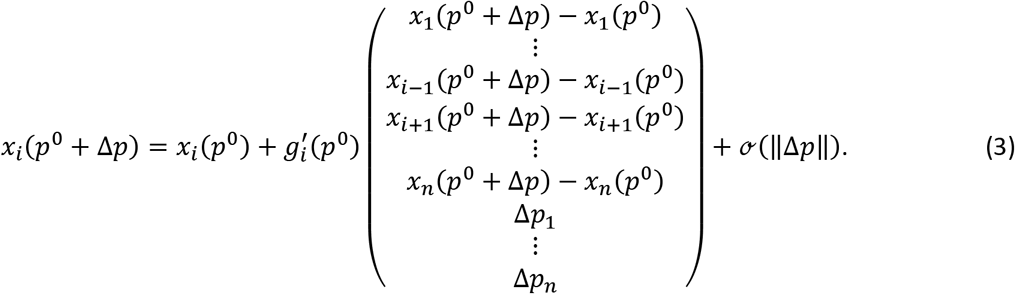

Dividing each side by *x_i_*(*p*^0^), Eq. (3) can be rewritten

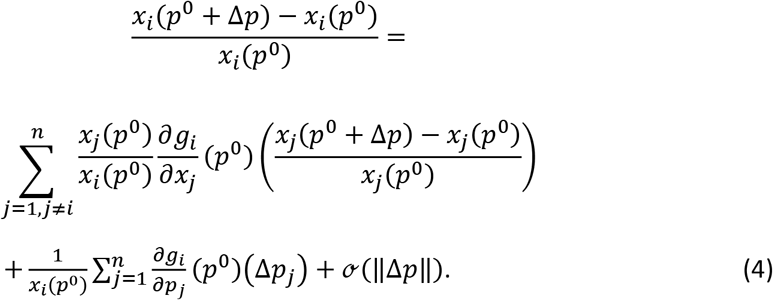

Since parameter *p_j_* influences module *j* only, 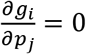 if *j ≠ i*. Moreover, *g_i_*(*x_j≠i_,p*) = *x_i_*(*x_j≠i_,p*) in *V_i_ × W_i_*, and if we denote

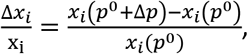

and

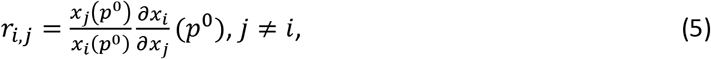

then

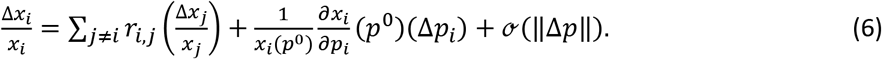

We next consider elementary perturbations *q_k_, k* ∈ {1,…,*n*}, which only perturb module *k*, i.e., the parameter *p_k_*. Neglecting the second-order term 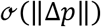 and writing

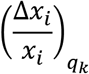

the relative difference in module *i* activity upon Δ*p_k_* change induced by perturbation *q_k_*, we find

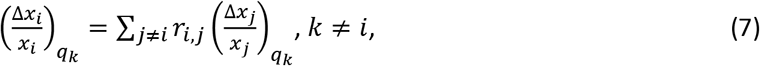

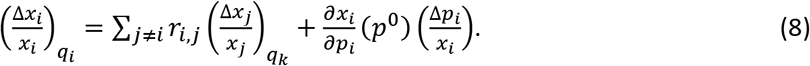

By defining *r_i,i_* = −1, we can write Eqs (7) and (8) in matrix form:

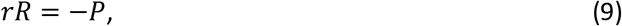

where *R* is the matrix that contains the experimentally observed relative activity changes 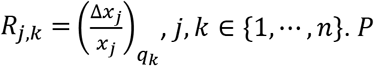 is a diagonal matrix with 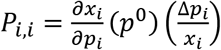, *i* ∈ {1,…,*n*}. The system (9) can be solved in two steps [1]. Firstly, *r* = −*PR*^−1^ and because *r_i,i_* = −1, we have *P_i,i_*(*R*^−1^)_i,i_ = 1 and 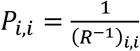. Secondly,

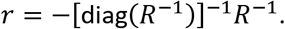

The elements of *R* are defined by 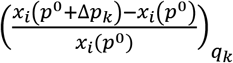 but as previously suggested [1], we preferred to estimate this quantity by

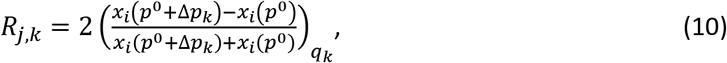

which avoids divisions by 0 and is numerically more stable.

Finally, from Eq. (5), we see that *r_i,j_* contains the searched coupling between MRA modules: direct action of *j* on *i* normalized by the ratio *X_j_/x_i_*. Similarly, *P_i,i_* measures the relative effect of *q_i_* on *x_i_*. We call it *q_i_* magnitude. The implicit function theorem provides analytical expressions for *g_i_*′ in terms of *f* partial derivatives, but since *f* is generally unknown, we did not use them. In his seminal work, Kholodenko made additional hypotheses to show that *r* contains coupling information, which is not necessary with our derivation. To be rigorous, one should ultimately restrict the model to neighborhoods included in all the *V_i_*’s, *W_i_*’s, and *U_i_*’s.

MRA models have been largely used for their inference capabilities [4]. Let us define a multiple perturbation *q* to be the linear combination of elementary perturbations *q_k_*. For instance, a perturbation on modules *i* and *j* with the same individual magnitudes would be coded by a column vector *c* with 1’s at positions *í* and *j* and 0’s elsewhere. From Eq. (9), we compute

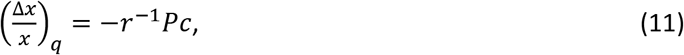

with 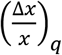 the column vector containing the inferred relative changes on each module activity.

Denoting Δ*p* the parameter changes induced by *q*, individual module activities are given by

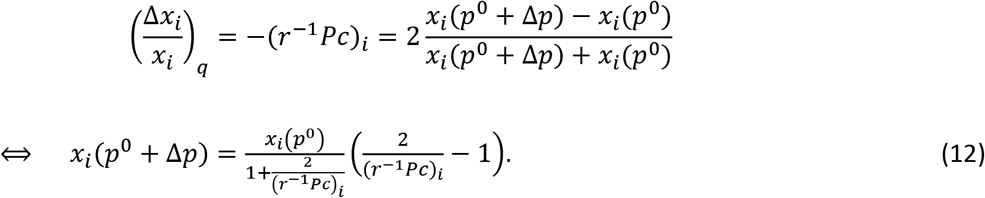

In case elementary perturbations contribute for different amounts to *q*, the vector *c* contains *q_k_*’s relative weights. In every case, linearity between the perturbation strength and its impact on *p* is assumed.

Confidence intervals (CI) around model parameters are estimated by a bootstrap procedure [17]. We considered experimental designs with biological and technical replicates. We average the technical replicates to obtain one activity value *per* biological replicate. The latter are again averaged to compute the *R* matrix according to Eq. (10). From the biological replicates, we estimate the variance of each *x_i_* employing an estimator optimized for a small sample size from Statistical Process Control theory [18,19]. Finally, a Gaussian distribution is assumed and 10^6^ *R* matrices are generated, which are submitted to MRA computations. The 95% CI is obtained from the 2.5^th^ and 97.5^th^ percentiles. In case 0 is not included in the CI, the MRA parameter is deemed significant and marked by an asterisk in the figures.

Inferences obtained from Eq. (12) were also complemented by the estimation of CIs following the principles above.

### Transcriptional data

ERβ and RARβ expression could not be quantified in MELN cells. We hence learned networks involving an ERα module, which transcriptional activity was reported by the ERE/luciferase construct. That is, luciferase mRNA abundance measured ERα activity. ERα mRNA abundance would combine ligand-bound and free amounts of the receptor, but only the ligand-bound ones matter in the model. We did not try to distinguish between RARα and RARγ. We estimated their combined transcriptional activity by the mRNA abundance of the *HOXA5* gene and the corresponding MRA module was named RARs. NRIP1 and LCoR activity were determined by their gene mRNA abundance. Since MELN cells are BC cells, we considered the E2-, RA-, or E2+RA-stimulated conditions as basal. That is, perturbations at ERα and RARs were negative (switch to ethanol). Perturbations at NRIP1 and LCoR were achieved by siRNAs, i.e., they were also negative.

### The ERα-NRIP1-LCoR network

In an unstimulated condition (no E2), it is known that NRIP1 expression induces LCoR expression [10]. We started by assessing this coupling under the E2 basal condition and found similar coupling (Fig. 2A). We also observed negative coupling from LCoR to NRIP1, which is logical since NRIP1 is a direct target of E2-bound ERα and LCoR one of its corepressor. We next inferred the ERα-NRIP1-LCoR network under E2 (Fig. 2B). We could observe the known induction of NRIP1 by ERα with negative feedback [6]. We also reconstituted the known inhibition of ERα by LCoR [9]. Interestingly, the induction of LCoR upon NRIP1 expression observed in Fig. 2A became a double inhibition via ERα in Fig. 2B. This makes sense since there is no transcriptional control by NRIP1 alone, it can only modulate ERα activity. Perturbation magnitudes are in Fig. 2C. Finally, we assessed the validity of the inferred network by checking its predictive power. From Fig. 2D, we note a reasonable fidelity of the model and that the relative errors are commensurate with the CI sizes, i.e., with data variability.

**Fig. 2.**
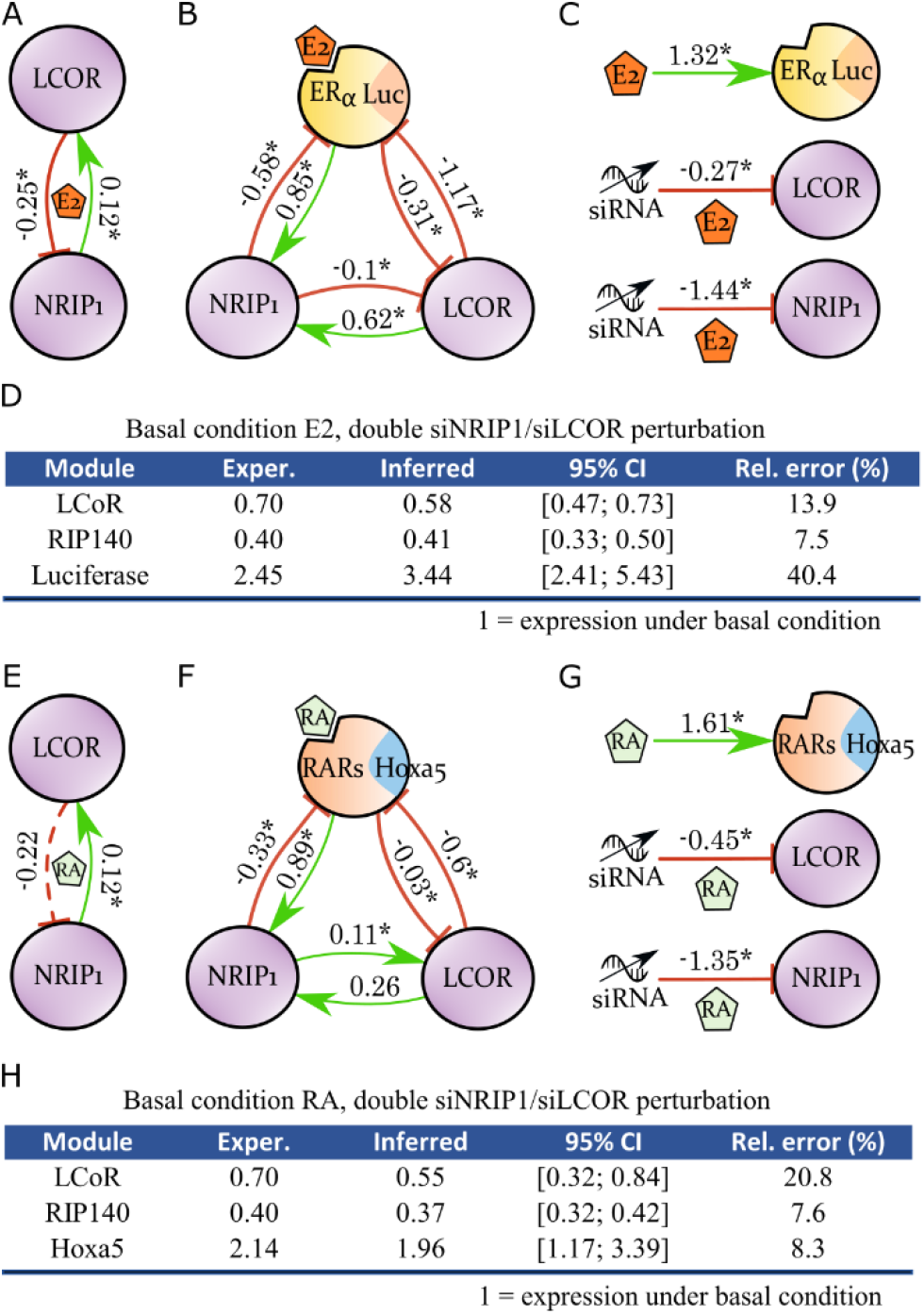
ER and RAR separated networks. **A.** Coupling between the two corepressors under E2 condition. A 95% CI for each model parameter was estimated and the parameter marked by an asterisk provided 0 was not included (nonzero with 5% significance). **B.** The ERα-NRIP1-LCoR transcriptional network. **C.** ERα, NRIP1, and LCoR perturbation magnitudes. **D.** Inference of gene expression under the dual siNRIP1 and siLCoR perturbation. **E.** Similar to A but under RA condition. **F.** The RARs-NRIP1-LCoR network. **G.** Similar to C. **H.** Similar to D.

### The RARs-NRIPl-LCoR network

Our next endeavor was to build a RARs-NRIP1-LCoR network before switching to the full network with both NRs. NRIP1 and LCoR coupling under RA stimulation (Fig. 2E) remained similar to its state under E2. That was expected since these two corepressors are used by several NRs. In Fig. 2F, we reconstituted the induction of NRIP1 expression by RAR as well as the inhibition of RAR expression by NRIP1 [11]. The inhibition of RAR by LCoR was also known [9]. Coupling between LCoR and NRIP1 is essentially similar to Fig. 2B since the NRIP1-to-LCoR arrows featured weak coupling. NRIP1 perturbation magnitude remained close, but LCoR perturbation changed 2-fold (Fig. 2G) although the same siRNAs were used. Inferences (Fig.2H) also supported the accuracy of the model.

### The full ERα-RARs-NRIP1-LCoR network

We followed the same approach as above to construct a full model of ER-RAR crosstalk (Fig. 3A). Perturbation magnitudes (Fig. 3B) were in the same range as before under the new dual E2 and RA basal condition. Values for NRIP1 and LCoR were closer to the RAR-NRIP1-LCoR network. No literature reports coupling with the corepressors NRIP1 and LCoR under this particular condition. Only the crosstalk between RAR and ER mentioned in the introduction is known [15,16]. We hence first challenged the model by testing its predictive accuracy (Fig. 3C), which was again satisfying.

**Fig. 3.**
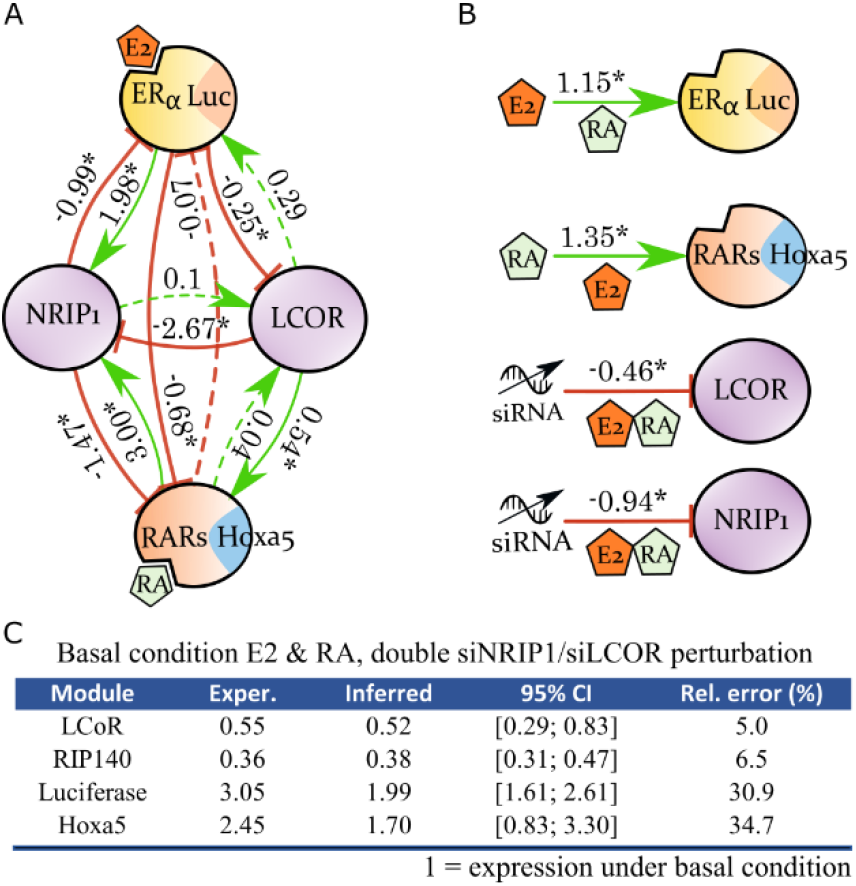
The ERα-RARs-NRIP1-LCoR network. **A.** MRA network model. **B.** Perturbation magnitudes under the dual E2 and RA stimulation. **C.** Inferred activity of the modules upon double siRNA inhibition of NRIP1 and LCoR.

Interestingly, cross inhibition of ER and RAR signaling acted along two paths. The model shows direct inhibition of ER transcriptional activity by the RAR module, which was described in the literature [15,16]. Reciprocal inhibition was suggested but not significant in our data. In agreement with our hypothesis, we found a parallel crosstalk mechanism through the induction of NRIP1 expression, which could subsequently repress both RARs and ERα. MRA modeling thus supported the coexistence of the two phenomena. Although LCoR reversed action on NRIP1 compared to the E2 and RA independent conditions might counterbalance cross inhibition of the two NRs, the strengths of coupling on the model edges and the much-attenuated induction of LCoR by NRIP1 suggested that it was not the case.

### MRA models from RNA-seq data

Since MRA relies on module activity relative change (Eqs. (7-8)), absolute quantitation is not necessary. We thus computed an RARs-NRIP1-LCoR MRA models as HOXA5, NRIP1, and LCoR mRNA abundances were available in our RNA-seq data (Fig. 4A). Comparing with the qPCR-based model in Fig. 2F, we notice that all the significant edges of Fig. 2F plus NRIP1-to-LCoR were recovered. The only change is very weak coupling LCoR-to-RARs (−0.03) that became slightly positive (0.13) with RNA-seq data. CIs were not determined for RNA-seq data since only two replicates were available.

**Fig. 4.**
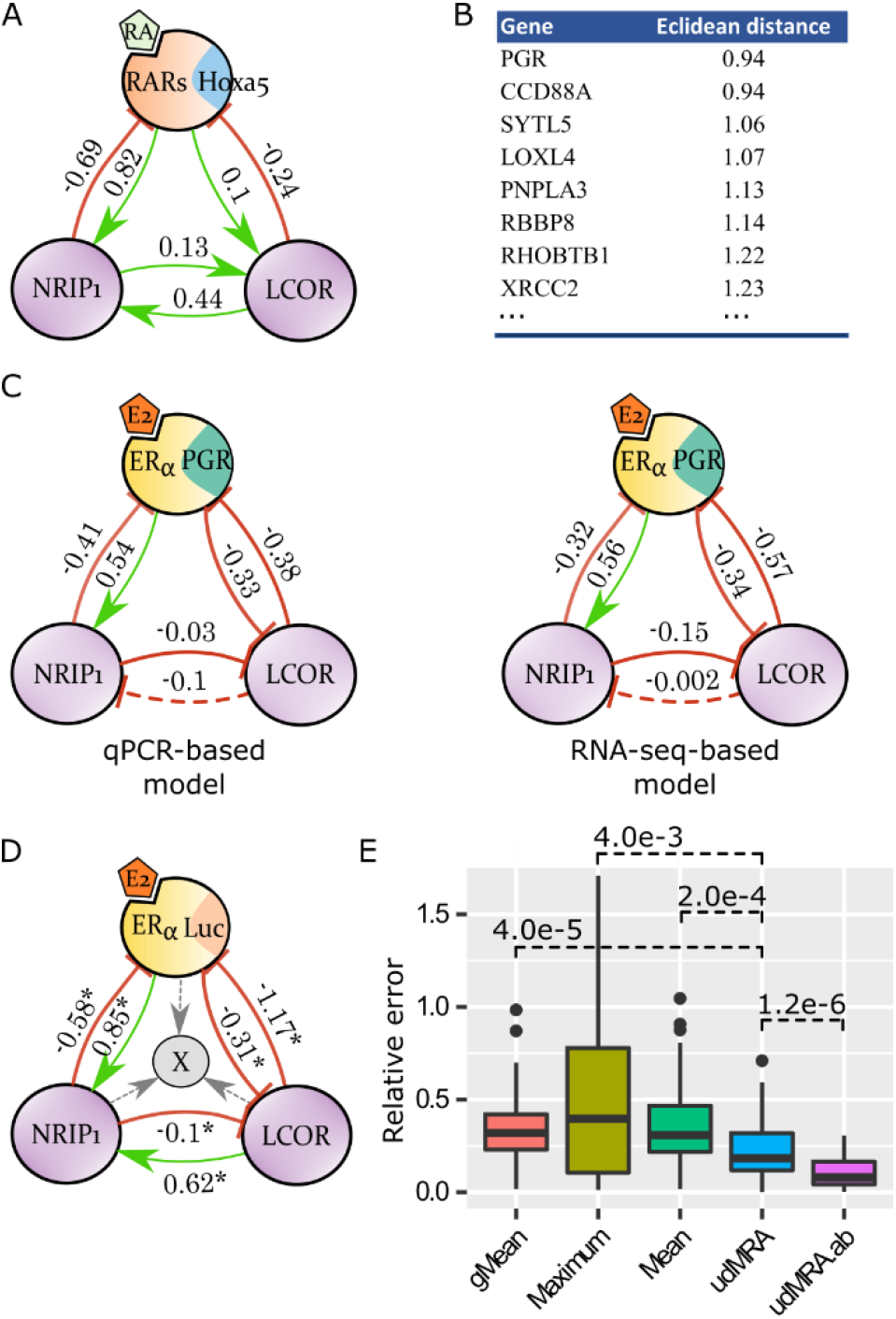
Genomic-scale inferences. **A.** RARs-NRIP1-LCoR model trained from RNA-seq data. **B.** Eight closest replacement genes for ERE-Luc in the ERα-NRIP1-LCoR model. **C.** ERα-NRIP1-LCoR models with ERE-Luc replaced by PGR, trained from qPCR and RNA-seq data. **D.** Principle of unidirectional MRA. **E.** Accuracy of unidirectional MRA inference (udMRA & udMRA.ab) under the E2 condition with double siNRIP1/siLCoR perturbation versus simple predictors (mean, geometric mean (gMean), and maximum of the two siRNAs).

The results above indicated that MRA could be applied to RNA-seq data. We, therefore, decided to exploit this opportunity by performing a new type of investigation. We used MRA to find a gene that would function as ERα transcription reporter, and would thus replace the ERE-Luc construct. Existing ChIP-seq data [20] intersected with our RNA-seq data allowed us to identify 884 genes targeted by ERα and E2-regulated (edgeR analysis, P-value<0.01, fold-change>2). We hence computed 884 MRA models with siNRIP1 and siLCoR RNA-seq data, replacing ERE-Luc by each of those genes successively. The genes with closest Euclidean distances between their model coupling parameters (the *r_i,j_* matrix) and those of the original Fig. 2B qPCR model are listed in Fig. 4B. We decided to test PGR and measured its expression by qPCR. The qPCR- and RNA-seq-based models are featured in Fig. 4C. They accurately reproduced the original model of Fig. 2B and were very similar to each other, thereby further validating the use of RNA-seq data for MRA model training.

### Unidirectional MRA on a genome-scale

We reasoned that the ERα-NRIP1-LCoR MRA model might provide means of predicting E2-regulated gene expression. We hence introduced a modified ERα-NRIP1-LCoR MRA model with one additional module that cannot influence the other modules (Fig. 4D). The gray unidirectional arrows in Fig. 4D represent the coupling between NRIP1, LCoR, the ERα module, and the added gene denoted by X. This coupling can be learned in the E2 basal condition by applying elementary perturbations as above. This must be repeated for each gene X considered. Gene X mRNA abundance is an *n* + 1^th^ module and, by hypothesis, *r*_*i,n*+1_ = 0, ∀*i* ∈ {1,…, *n*}, since no return arrows exist. From Eq. (7), we can compute *r*_*n*+1,*j*_, ∀*j* ∈ {1,…,*n*}, by solving the system

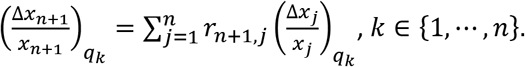

The performance of this new type of MRA model (udMRA) was assessed by its ability to predict module *n* + 1 activity under the dual siNRIP1/siLCoR condition, as we did above for the other models. To avoid trivially successful predictions on genes that would not vary, we limited the benchmark to the 884 genes above that were also significantly regulated upon siNRIP1 or siLCoR in the E2-stimulated condition. That left us with 60 genes. In Fig. 4E, we report the relative errors observed applying udMRA and comparing with naïve predictions. udMRA yielded significantly better estimates of the added module activity.

One could wonder whether perturbation magnitudes during double siRNA interference on the same biological system remain identical. That is, whether filling the vector *c* in Eq. (11) with 1’s at the perturbed module indices (what we did so far) is the best option. Eq. (11) is written such that we can try different values. We searched for optimal coefficients *a* and *b* applied to siNRIP1 and siLCoR perturbations (at the corresponding indices in vector *c*), such that prediction errors of Luciferase, NRIP1, and LCoR expression (as in Fig. 2D) would be minimal. We found *a* = 1, and *b* = 0.4. Then, we used those coefficients in the udMRA model to try to predict the expression of the 61 benchmark genes more accurately. Indeed, we see in Fig. 4E that this new model called udMRA.ab achieved much better accuracy.

### aiMeRA usage

The R package was designed to be generally applicable; it relies on the formulae presented here. It is able to work with any quantitative input, including biological and technical replicates. We included functionality to facilitate the definition of MRA model topologies (Fig. 5A). Model construction only involves the execution of a few generic R functions and network plots can be generated within R directly (Fig. 5B). It is also possible to export such graphs in the graphML format for loading into Cytoscape [21]. More details are provided in the package documentation. aiMeRA is available from GitHub; submission to Bioconductor is pending.

**Fig. 5.**
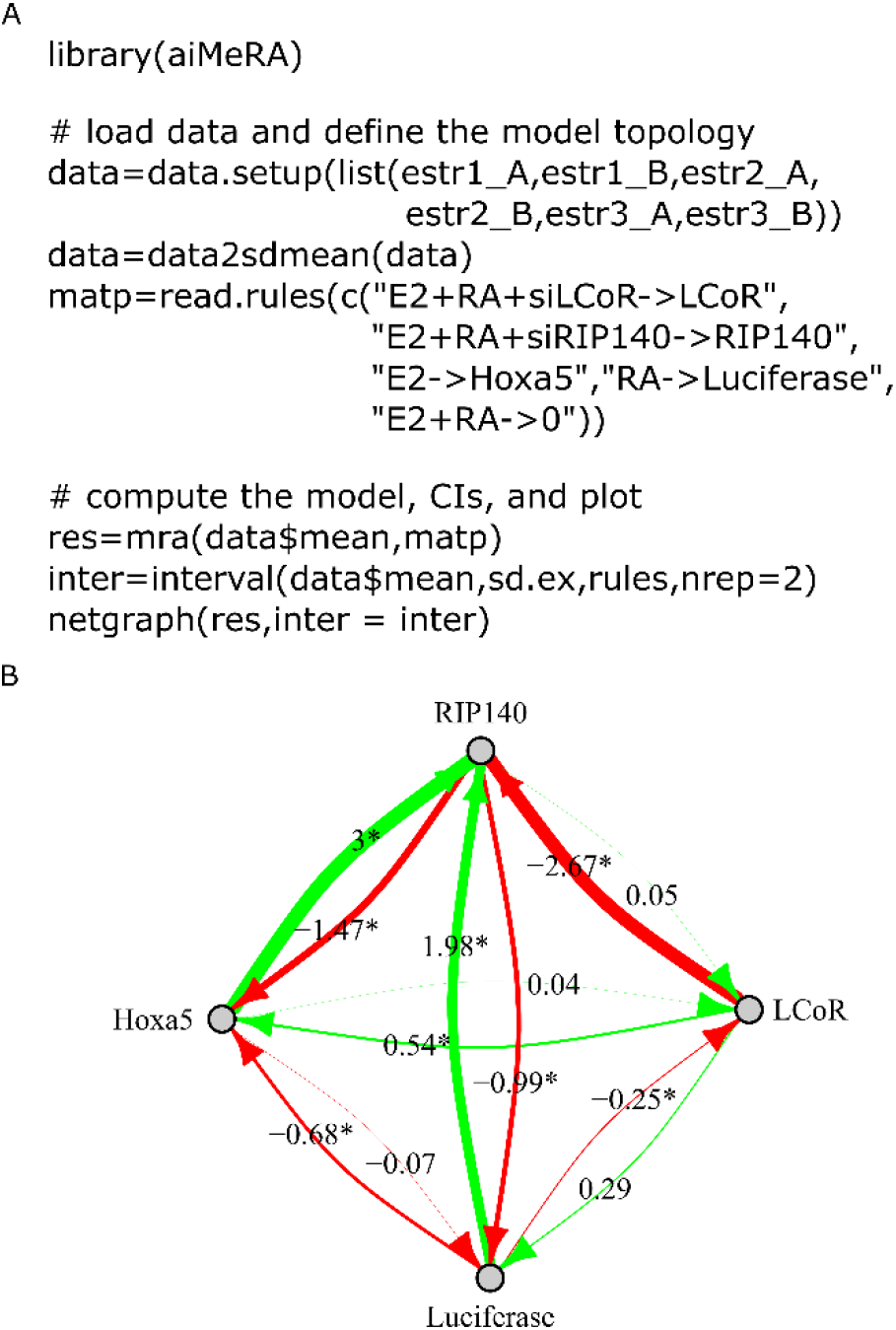
The aiMeRA R package. **A.** Example R code to load data, prepare them and compute the model. Note that NRIP1 was called after its common alternative name RIP140. Basal condition is E2 plus RA and we see that LCoR perturbation is defined as E2+RA+siLCoR. Same logic for RIP140 (=NRIP1). Perturbation on the HOXA5 module reporting RARs activity is defined as E2, i.e., loss of RA stimulation. Etc. **B.** Plot of an MRA model in R using the igraph library.

## Discussion

MRA modeling [1,2] is a widely used technique to learn the coupling between the modules of a biological system from perturbation data sets (Fig. 1). We introduced a new mathematical derivation of the method and implemented a generic R package called aiMeRA (Fig. 5). Application thereof was illustrated on an unpublished data set combining specific qPCR and broad RNA-seq data to explore crosstalk between ER and RAR, two important NRs involved in several tumors such as BC. Data analysis reconstituted some known interactions (Fig. 2) and supported a novel hypothesis that reciprocal negative coupling could be mediated by shared corepressors (Fig. 3).

We showed that MRA transcriptional models trained from RNA-seq data are close to those trained from qPCR (Fig. 4A-C). Which lead us to introduce an innovative application of MRA to probe genes genome-wide, searching for replacement reporters of module activity. That is, new genes that could be functionally related to an MRA module. In particular, we found that the progesterone receptor gene (*PGR*) reported on ligand-bound ERα transcriptional activity accurately. That observation indicates the potential value of this new use of MRA models since PGR is a widely used reporter of estrogen activity in BC in the clinic.

We additionally investigated the possibility to build hybrid MRA models (udMRA & udMRA.ab) including unidirectional coupling to add modules that were not perturbed in the training data set (Fig. 4D). In the context of the application presented in this report, i.e., the transcriptional activity of a system of transcription factors, we showed that the ERα-RARs-NRIP1-LCoR udMRA and udMRA.ab models could outperform naïve predictors (Fig. 4E). Other biological systems might be amenable to such modified models in the absence of strong back-coupling. The aiMeRA package methods support both RNA-seq data and udMRA models.

## Methods

### Cell culture and perturbation experiments

We used MELN cells, an MCF7-derived cell line stably transfected with the estrogen-responsive luciferase reporter gene ERE-βGlob-Luc-SV-Neo [22]. The cell line was authenticated by short tandem repeat profiling and tested for mycoplasma contamination.

MELN cells were cultured in phenol red-free Dulbecco’s modified Eagle medium (Gibco) containing 5% dextran-charcoal treated FCS (Invitrogen) and antibiotics (Gibco). Perturbations at NRIP1 and LCoR were obtained by siRNAs that were transfected using Interferrin (Polyplus). Perturbations at ERα and RARs were induced by their respective natural ligands: the hormones estrogen (17β-estradiol or E2 for short) and all-trans retinoic acid (RA).

MELN cells were obtained in the following conditions: basal (untreated), E2, RA, E2+RA, siNRIP1, siLCOR, siNRIP1+siLCOR, E2+siNRIP1, E2+siLCOR, E2+siNRIP1+siLCOR, RA+siNRIP1, RA+siLCOR, RA+siNRIP1+siLCOR, E2+RA+siNRIP1, E2+RA+siLCOR, and E2+RA+siNRIP1+siLCOR. These experiments were realized in triplicates. Cells were harvested after 18 hours of culture. E2-treated cells received 100nM E2, RA-treated cells 10 uM RA, and untreated cells ethanol. Validations of the response to E2 and siRNA interference are in Suppl. Fig. 1.

### mRNA quantification

RNA was isolated using the Zymo Research kit (Zymo Research) and reverse transcription (RT)-qPCR assays were done using qScript (VWR) according to the manufacturer’s protocol. Transcripts were quantified using SensiFAST SYBR (BioLine) on an LC480 instrument. The nucleotide sequences of the primers used for real-time PCR were:

RIP140-f (5’- AATGTGCACTTGAGCCATGATG -3’),
RIP140-r (5’- TCGGACACTGGTAAGGCAGG -3’),
LCoR-f (5’- GAACCTAGCGAACAAGACGGTG -3’),
LCoR-r (5’- TGGAGAGTGGCTCAGGGAAGT -3’),
Luciferase-f (5’- CTCACTGAGACTACATCAGC -3’),
Luciferase-r (5’- TCCAGATCCACAACCTTCGC -3’),
HOXA5-f (5’- GCGCAAGCTGCACATAAGTC -3’),
HOXA5-r (5’- GAACTCCTTCTCCAGCTCCA -3’),
ERα-f (5’- TGGAGATCTTCGACATGCTG -3’),
ERα-r (5’- TCCAGAGACTTCAGGGTGCT -3’),
RARα-f (5’- GGATATAGCACACCATCCCC -3’),
RARα-r (5’- TTGTAGATGCGGGGTAGAGG -3’),
PGR-f (5’- CGCGCTCTACCCTGCACTC-3’),
PGR-r (5’-TGAATCCGGCCTCAGGTAGTT-3’).
(RT)-qPCR data are available from the R package.

### mRNA sequencing

For two of the triplicates, in each condition, RNA was extracted as above described. Libraries were prepared with Illumina TruSeq kit and submitted to NextSeq500 sequencing (1×75bp/40M reads). The first 13 and last 7 bps were cut by an in-house Perl script to eliminate compositional bias. Cut reads were submitted to sickle to eliminate remaining low-quality regions. Alignments were performed against the human genome (hg38) with TopHat v2.10 [23] and read counts extracted with HTSeq-Count [24]. The read count matrix was normalized with edgeR [25] TMM algorithm. Data are available from GEO under GSE143956.

### aiMeRA library implementation

We implemented the MRA method according to the mathematical formulation above as an R library. (RT)-qPCR data of this project were embedded in the R library for convenience and to provide an example. We also included the data used in the MRA original paper [1] such that users can check that our code gives the same results as those reported in the latter publication.

## Acknowledgments

We thank Simon Cabello-Aguilar and Meriem Mekedem for useful discussions during the development of the project. JC was supported by the Agence Nationale de la Recherche (grant ANR-16-CE17-0002-02), the Groupement des entreprises françaises dans la lutte contre le cancer (GEFLUC), and the Fondation ARC (PJA 20151203332).

## Supporting information captions

**Suppl. Fig. 1. A.** MELN cells were transfected with either a siRIP140 (siNRIP1), a siLCoR or a combination of the two siRNAs. RIP140 and LCoR mRNA levels were quantified by real time PCR. Results are corrected to 28S mRNA and normalized to cells transfected with the control siRNA. **B.** MELN cells were transfected as described in A and treated with estradiol (10^−7^ M) when indicated. Luciferase mRNA expression is quantified as in A.

## References

1. Kholodenko BN, Kiyatkin A, Bruggeman FJ, Sontag E, Westerhoff HV, Hoek JB. Untangling the wires: a strategy to trace functional interactions in signaling and gene networks. Proc Natl Acad Sci U S A. 2002;99: 12841–6. doi:10.1073/pnas.192442699

2. Santra T, Rukhlenko O, Zhernovkov V, Kholodenko BN. Reconstructing static and dynamic models of signaling pathways using Modular Response Analysis. Current Opinion in Systems Biology. 2018;9: 11–21. doi:10.1016/j.coisb.2018.02.003

3. Dorel M, Klinger B, Gross T, Sieber A, Prahallad A, Bosdriesz E, et al. Modelling signalling networks from perturbation data. Bioinformatics. 2018;34: 4079–4086. doi:10.1093/bioinformatics/bty473

4. Klinger B, Sieber A, Fritsche-Guenther R, Witzel F, Berry L, Schumacher D, et al. Network quantification of EGFR signaling unveils potential for targeted combination therapy. Mol Syst Biol. 2013;9: 673. doi:10.1038/msb.2013.29

5. Heldring N, Pike A, Andersson S, Matthews J, Cheng G, Hartman J, et al. Estrogen receptors: how do they signal and what are their targets. Physiol Rev. 2007;87: 905–931. doi:10.1152/physrev.00026.2006

6. Cavaillès V, Dauvois S, L’Horset F, Lopez G, Hoare S, Kushner PJ, et al. Nuclear factor RIP140 modulates transcriptional activation by the estrogen receptor. EMBO J. 1995;14: 3741–3751.

7. Docquier A, Garcia A, Savatier J, Boulahtouf A, Bonnet S, Bellet V, et al. Negative regulation of estrogen signaling by ERβ and RIP140 in ovarian cancer cells. Mol Endocrinol. 2013;27: 1429–1441. doi:10.1210/me.2012-1351

8. Sixou S, Müller K, Jalaguier S, Kuhn C, Harbeck N, Mayr D, et al. Importance of RIP140 and LCoR Sub-Cellular Localization for Their Association With Breast Cancer Aggressiveness and Patient Survival. Transl Oncol. 2018;11: 1090–1096. doi:10.1016/j.tranon.2018.06.006

9. Fernandes I, Bastien Y, Wai T, Nygard K, Lin R, Cormier O, et al. Ligand-Dependent Nuclear Receptor Corepressor LCoR Functions by Histone Deacetylase-Dependent and -Independent Mechanisms. Molecular Cell. 2003;11: 139–150. doi:10.1016/S1097-2765(03)00014-5

10. Jalaguier S, Teyssier C, Nait Achour T, Lucas A, Bonnet S, Rodriguez C, et al. Complex regulation of LCoR signaling in breast cancer cells. Oncogene. 2017;36: 4790–4801. doi:10.1038/onc.2017.97

11. White KA, Yore MM, Warburton SL, Vaseva AV, Rieder E, Freemantle SJ, et al. Negative Feedback at the Level of Nuclear Receptor Coregulation SELF-LIMITATION OF RETINOID SIGNALING BY RIP140. J Biol Chem. 2003;278: 43889–43892. doi:10.1074/jbc.C300374200

12. White KA, Yore MM, Deng D, Spinella MJ. Limiting effects of RIP140 in estrogen signaling: potential mediation of anti-estrogenic effects of retinoic acid. J Biol Chem. 2005;280: 7829–7835. doi:10.1074/jbc.M412707200

13. Rousseau C, Pettersson F, Couture MC, Paquin A, Galipeau J, Mader S, et al. The N-terminal of the estrogen receptor (ERalpha) mediates transcriptional cross-talk with the retinoic acid receptor in human breast cancer cells. J Steroid Biochem Mol Biol. 2003;86: 1–14. doi:10.1016/s0960-0760(03)00255-3

14. Laganière J, Deblois G, Giguère V. Functional genomics identifies a mechanism for estrogen activation of the retinoic acid receptor alpha1 gene in breast cancer cells. Mol Endocrinol. 2005;19: 1584–1592. doi:10.1210/me.2005-0040

15. Hua S, Kittler R, White KP. Genomic antagonism between retinoic acid and estrogen signaling in breast cancer. Cell. 2009;137: 1259–1271. doi:10.1016/j.cell.2009.04.043

16. Ross-Innes CS, Stark R, Holmes KA, Schmidt D, Spyrou C, Russell R, et al. Cooperative interaction between retinoic acid receptor-alpha and estrogen receptor in breast cancer. Genes Dev. 2010;24: 171–182. doi:10.1101/gad.552910

17. Davison AC, Hinkley DV. Bootstrap Methods and their Applications. Cambridge University Press; 1997. Available: https://doi.org/10.1017/CBO9780511802843

18. Wheeler DJ, Chambers DS. Understanding Statistical Process Control. 2nd ed. SPC Press-Inc.; 1992.

19. Harter HL. Tables of Range and Studentized Range. Ann Math Statist. 1960;31: 1122–1147. doi:10.1214/aoms/1177705684

20. Madak-Erdogan Z, Charn T-H, Jiang Y, Liu ET, Katzenellenbogen JA, Katzenellenbogen BS. Integrative genomics of gene and metabolic regulation by estrogen receptors α and β, and their coregulators. Mol Syst Biol. 2013;9: 676. doi:10.1038/msb.2013.28

21. Smoot ME, Ono K, Ruscheinski J, Wang PL, Ideker T. Cytoscape 2.8: new features for data integration and network visualization. Bioinformatics. 2011;27: 431–2. doi:10.1093/bioinformatics/btq675

22. Balaguer P, Boussioux AM, Demirpence E, Nicolas JC. Reporter cell lines are useful tools for monitoring biological activity of nuclear receptor ligands. Luminescence. 2001;16: 153–158. doi:10.1002/bio.630

23. Kim D, Pertea G, Trapnell C, Pimentel H, Kelley R, Salzberg SL. TopHat2: accurate alignment of transcriptomes in the presence of insertions, deletions and gene fusions. Genome Biology. 2013;14: R36. doi:10.1186/gb-2013-14-4-r36

24. Anders S, Pyl PT, Huber W. HTSeq--a Python framework to work with high-throughput sequencing data. Bioinformatics. 2015;31: 166–169. doi:10.1093/bioinformatics/btu638

25. McCarthy DJ, Chen Y, Smyth GK. Differential expression analysis of multifactor RNA-Seq experiments with respect to biological variation. Nucleic Acids Res. 2012;40: 4288–4297. doi:10.1093/nar/gks042

